# Resilience, Invariability, and Ecological Stability across Levels of Organization

**DOI:** 10.1101/085852

**Authors:** Bart Haegeman, Jean-François Arnoldi, Shaopeng Wang, Claire de Mazancourt, José M. Montoya, Michel Loreau

## Abstract

Ecological stability is a bewildering broad concept. The most common stability measures are asymptotic resilience, widely used in theoretical studies, and measures based on temporal variability, commonly used in empirical studies. We construct measures of invariability, defined as the inverse of variability, that can be directly compared with asymptotic resilience. We show that asymptotic resilience behaves like the invariability of the most variable species, which is often a rare species close to its extinction boundary. Therefore, asymptotic resilience displays complete loss of stability with changes in community composition. In contrast, mean population invariability and ecosystem invariability are insensitive to rare species and quantify stability consistently whether details of species composition are considered or not. Invariability provides a consistent framework to predict diversity-stability relationships that agree with empirical data at population and ecosystem levels. Our findings can enhance the dialogue between theoretical and empirical stability studies.

## Introduction

How do ecosystems respond to perturbations? Although this is the driving question of the decades-old field of ecological stability, the insights gained are still fragmentary (May, 1973; McCann, 2000; Ives & Carpenter, 2007). The lack of integrative results is at least partly due to the bewildering extent of the concept of ecological stability (Pimm, 1984; Grimm & Wissel, 1997; Donohue *et al.*, 2013). By combining a particular type of perturbation (change in an environmental parameter, biomass addition or removal, species extinction or invasion; of short, long or lasting duration; of weak or strong intensity; affecting one, several or all populations, …) with a particular type of system response (immediate, short- or long-term; at local or larger spatial scale; at population or community level, …), a multitude of stability notions and measures can be defined and have indeed been used in the literature. While this conceptual diversity might be seen as a reflection of the complexity of the real world, it is severely hampering the progress of the field. Without systematic links between stability notions, it is impossible to integrate the results of individual studies into a more comprehensive stability theory (Donohue *et al.*, 2013; Arnoldi *et al.*, 2016).

Particularly problematic is the disparity in the way stability is quantified in theoretical and empirical studies (Donohue *et al.*, 2016). The majority of theoretical studies focuses on the mathematical notion of asymptotic stability, i.e., whether the system returns after a small pulse perturbation to its pre-perturbation state, which is most often an equilibrium point (May, 1973; Allesina & Tang, 2013; Coyte *et al.*, 2015). The associated quantitative stability measure, asymptotic resilience, measures the long-term rate of return to equilibrium (e.g., Neutel *et al.*, 2002; Rooney *et al.*, 2006; Thébault & Fontaine, 2010; Tang *et al.*, 2014). The larger this return rate, the more stable the system. To avoid confusion, it should be noted here that another ecological stability notion is also called resilience (Holling, 1973; Gunderson, 2000), which is however less often quantified in the theoretical literature. On the other hand, the most popular way of measuring stability in empirical studies is based on temporal variability, typically using the coefficient of variation of species or community biomass (Tilman *et al.*, 2006; Jiang & Pu, 2009; Campbell *et al.*, 2011; Gross *et al.*, 2014). Stability is then inversely related to variability, the idea being that more variable systems function in a less consistent way. As it is commonly believed that these two stability notions are intrinsically disconnected, theoretical and empirical approaches to ecological stability have become widely divergent.

Here we intend to narrow the gap between theoretical and empirical stability studies by examining the similarities and differences between asymptotic resilience and temporal variability. Counter to a widespread belief, we emphasize that these two stability notions are *not* fundamentally incomparable (Fig. 1). First, several empirical studies have estimated asymptotic resilience from time-series data that describe the response of an ecosystem to a natural or artificial pulse perturbation (e.g., Steiner *et al.*, 2006; Sibly *et al.*, 2007). Because the system progressively approaches the equilibrium, the asymptotic return rate is often hidden in the naturally occurring fluctuations around the equilibrium. The usual workaround consists in using the return rate at a shorter time as a proxy. Second, several theoretical studies have studied the properties of temporal variability (May, 1974; Ives *et al.*, 1999; Lehman & Tilman, 2000; Loreau & de Mazancourt, 2008). To this end, a persistent, stochastic perturbation is applied to a model ecosystem and the intensity of its stationary fluctuations is quantified. In this paper we take this approach a step further by introducing a variability-based stability measure, which we call invariability, that is directly comparable to asymptotic resilience. Our measure coincides with asymptotic resilience for the simplest systems, but deviates from it for more complex ones.

Thus, resilience and invariability describe the response of a system to different types of perturbations, i.e., a pulse and a persistent, stochastic perturbation, respectively. The correspondence between the two concepts can be made explicit, however, by noting that a persistent perturbation can be seen as being composed of a sequence of pulse perturbations. Hence, variability is a sum of short- and long-term responses to pulse perturbations (Arnoldi *et al.*, 2016). But the two stability notions also have fundamentally different properties, as can be seen from comparing the system’s response for different variables (Fig. 1). The asymptotic rate of return to equilibrium is the same for each species, and also for total biomass. Thus, asymptotic resilience can be interpreted as a rigid stability property. In particular, it does not change across levels of organization (population to ecosystem). In contrast, the variability of total biomass is typically smaller than that of individual species, as species fluctuations are averaged out when taking the sum of their biomass. Hence, variability can be used to distinguish between stability at the population and ecosystem levels, as has been done in empirical studies (Tilman, 1996; Jiang & Pu, 2009; Campbell *et al.*, 2011).

We provide novel insights into the relationship between asymptotic resilience and variability-based stability. We show that asymptotic resilience can be interpreted as an extreme invariability measure, namely the invariability of the most variable population. This suggests that asymptotic resilience should be considered as a population-level stability measure. Furthermore, we show that both the minimal population invariability and asymptotic resilience drop to zero each time a species is integrated into or lost from the community. This leads to repeated complete loss of stability along environmental gradients with species turnover, as is often the case in nature. We show that other, less extreme invariability measures do not lose stability when species composition changes. We introduce two such measures, one at the population level and one at the ecosystem level, and show that they quantify stability consistently across levels of organization.

Finally, we discuss the implications of these findings for the heated debate on the relationship between diversity and stability (McCann, 2000; Ives & Carpenter, 2007). For competitive interactions we show that theory predicts different relationships depending on the choice of the stability measure: asymptotic resilience and population-level invariability suggest negative diversity-stability relationships, while ecosystem-level invariability show a positive relationship, in agreement with empirical data (Jiang & Pu, 2009; Campbell *et al.*, 2011; Gross *et al.*, 2014). Thus, understanding the differences between asymptotic resilience and invariability sheds new light on the controversies surrounding the diversity-stability relationship.

## Measures of population and ecosystem stability

To introduce the various stability measures, consider an ecosystem model given by a set of differential equations that describe how the biomass *N_i_* of species *i* changes through time due to its interactions with other species and the environment. Assume that the model has an equilibrium point *N^∗^* (a vector of length *n*, where *n* is the number of species) and that the dynamics in the neighborhood of this equilibrium are described by community matrix *A* (components *A_ij_*, with *i* = 1*, …, n* and *j* = 1*, …, n*). Although the equilibrium assumption is not always appropriate for the study of ecological stability, it is widely adopted both in theoretical and empirical studies (see also Discussion).

**Figure 1:**
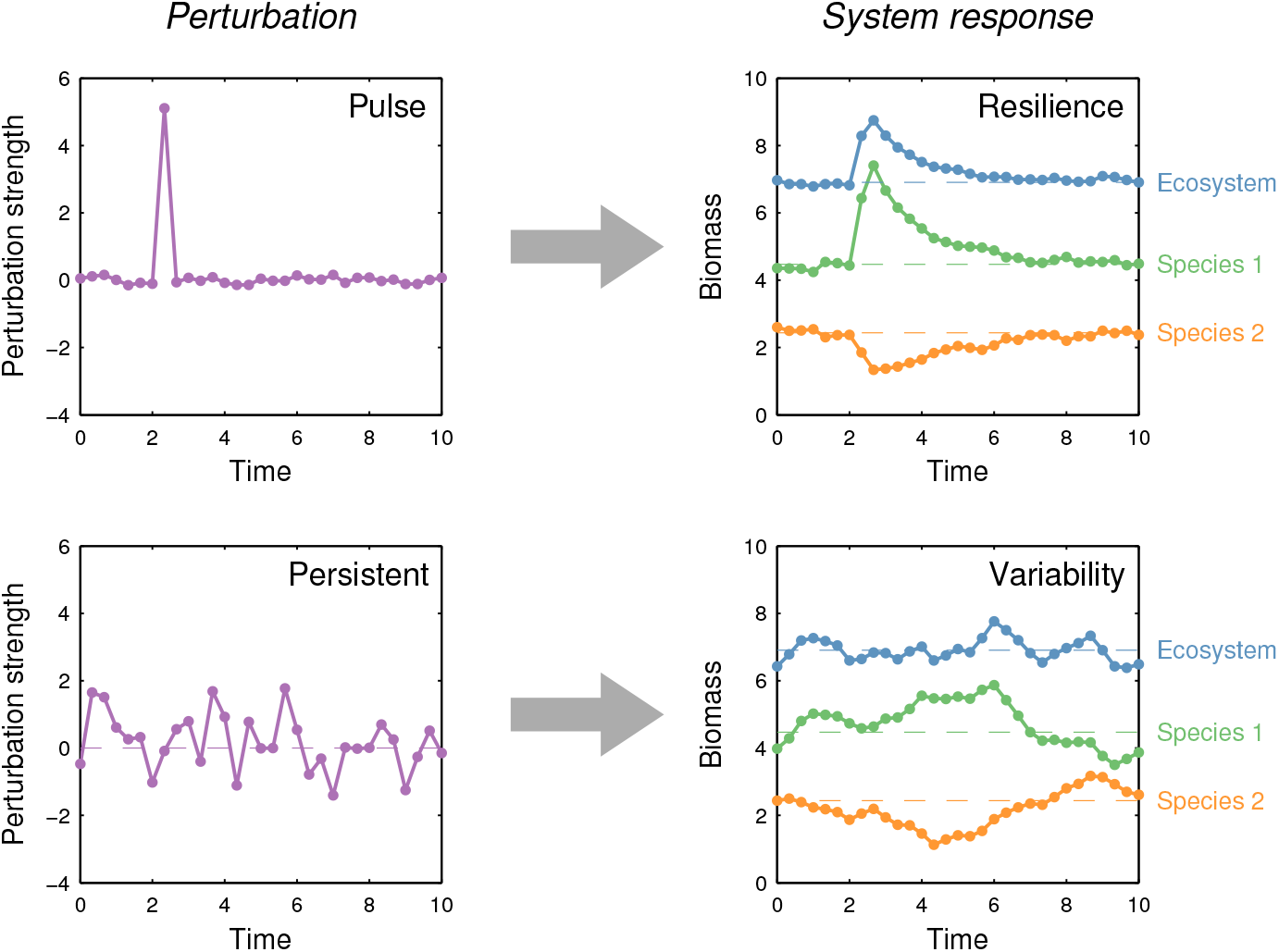
Resilience and variability are related stability measures, but they have different properties. Most stability measures are based on quantifying the system’s response to a perturbation. (Top) In case of a pulse perturbation, asymptotic resilience can be measured as the long-term rate of return to equilibrium. (Bottom) When the peturbation persists over time, temporal variability of system properties is used to quantify stability. The link between resilience and variability appears clearly when considering a persistent perturbation as a sequence of pulse perturbations. Nevertheless, resilience and variability have different properties. For example, the same long-term return rate is obtained from the time series of individual species biomass (green and orange) and of total biomass (blue). In contrast, variability of total biomass is typically smaller than species-level variability, especially when species fluctuate out of synchrony.

The linear dynamical system with community matrix *A* is stable if all its eigenvalues have negative real part. Asymptotic resilience *ℜ* is then related to the eigenvalue with the largest real part, also called the dominant eigenvalue *λ*_dom_,

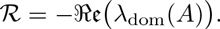

It is equal to the asymptotic rate of return to equilibrium after a pulse perturbation. As mentioned in the introduction, this rate is the same for almost any pulse perturbation and is typically independent of the dynamical variable (e.g., species or ecosystem biomass) through which the system response is observed (see Fig. 1 and Appendix S1 for more details). This is a remarkable property, which sets apart asymptotic resilience from other stability measures.

To construct stability measures based on temporal variability, we apply a stochastic white-noise perturbation to the deterministic linear dynamical system. For simplicity, we assume that perturbations are independent between species, and that they act on a per capita basis, as is typically the case with environmental stochasticity (Lande *et al.*, 2003). As a result, the perturbation acting on the deterministic system is characterized by the equilibrium abundance vector *N^∗^* and a parameter *σ*^2^, which measures the intensity of the environmental fluctuations (see Appendix S1).

In the stationary state the dynamical variables fluctuate with covariance matrix *C*, which can be computed from the community matrix *A*, the equilibrium vector *N^∗^* and the perturbation intensity *σ*^2^ (see Appendix S1 for details). Species biomass *N_i_* has variance Var(*N_i_*) = *C_ii_* and squared coefficient of variation CV^2^(*N_i_*) = *C_ii_/*(*N_i_ ^∗^*)^2^, which increase proportionally to *σ*^2^ (for sufficiently small *σ*; see Fig. S1). We eliminate the latter, trivial dependence by normalizing with respect to *σ*^2^. Hence, we obtain a measure of the variability of species *i*, CV^2^(*N_i_*)*/σ*^2^, and by taking its inverse, a measure of its stability or “invariability”,

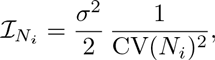

Invariability ℐ*_N_i__* is entirely determined by equilibrium *N^∗^* and community matrix *A*. It has units of 1/time as does asymptotic resilience *ℜ*, and coincides with asymptotic resilience for the simplest dynamical systems (see Appendix S1).

To construct population-level stability measures, we combine the invariabilities of individual populations into a single quantity. Here we introduce two such measures. First, we adopt the viewpoint that the most unstable component determines the stability of the system, and we consider the invariability of the most variable population,

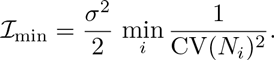

However, the species that has the minimum invariability ℐ_min_ need not be representative for the stability properties of the other species in the community. Alternatively, we construct a measure of population-level invariability by taking the average of the coefficients of variation weighted by species biomass (Thibaut & Connolly, 2013),

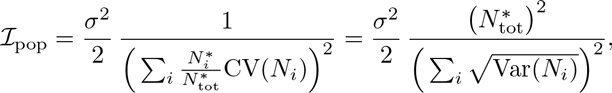

where 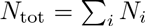 stands for the total biomass in the ecosystem. The resulting measure ℐ_pop_ is called population invariability and satisfies 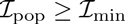. Note that a similar approach can be used to define population-level stability measures for a restricted set of species (e.g., species belonging to a specific trophic level).

To construct an ecosystem-level stability measure, we take the invariability of an aggregated ecosystem variable. Here we focus on total biomass 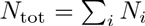. Its variance is equal to 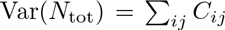 with *C* the covariance matrix of the dynamical variables. Then we define ecosystem invariability ℐ_eco_ as

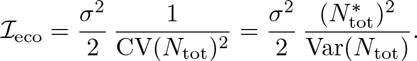

This measure satisfies the inequality ℐ_eco_ *≥ I*_pop_, which stems from the fact that population fluctuations are averaged out at the ecosystem level. When populations fluctuate more synchronously, ecosystem invariability decreases (relative to population invariability). In fact, the ratio of population and ecosystem invariability is a commonly used measure of ecosystem-wide synchrony (Loreau & de Mazancourt, 2008),

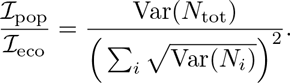

## Stability in the presence of species turnover

Resilience and invariability, which we have concretized as *ℜ*, ℐ_pop_ and ℐ_eco_, are widely used in the ecological literature. We now show that these measures lead to qualitatively different stability patterns, even in simple ecosystems. These differences are particularly striking in the presence of changes in species composition, as is typically the case when the stability of different ecosystems is compared. We illustrate this using a simple model, but our findings are quite general.

Consider a consumer-resource system in which three consumer species compete for two resources. Consumer species have different resource preferences; in the graphical theory of resource competition (Tilman, 1982), they have intersecting zero net growth isoclines, so that equilibrium coexistence depends on resource supply (see Fig. S2). We consider a gradient of resource supply, along which the equilibrium consumer community changes. Regions in which a single species dominates alternate with regions in which two species coexist (Fig. 2a). This type of compositional change patterns is common in studies of environmental gradients. To model an open ecosystem, we assume that each consumer species receives a small flow of immigrants. As a result, all species are present at equilibrium, but only the most competitive one or two (as predicted by resource competition theory) are common.

Asymptotic resilience *ℜ* drops to zero every time a consumer species is either lost from the community or able to invade the community (Fig. 2e). The fact that asymptotic resilience *ℜ* is very sensitive to changes in community composition is easily explained from a mathematical point of view: each compositional change corresponds to a transcritical bifurcation, in which an eigenvalue approaches zero. This eigenvalue is necessarily the dominant one, so that asymptotic resilience *ℜ* drops to zero. It is unclear, however, how to interpret this stability loss from an ecosystem viewpoint. At the bifurcations, community composition changes, but this change generally has only a minor effect on the ecosystem, as a rare species that disappears from the community does not affect the dominant species that largely determine the aggregated property of the ecosystem (e.g., total biomass). Hence, the stability loss predicted by asymptotic resilience is virtually irrelevant at the ecosystem level.

When white-noise perturbations are applied to consumer dynamics, the resulting fluctuations of population biomass are largest (largest variance) inside the coexistence regions due to the presence of a competitor (Fig. 2b). The invariability of a species is the lowest (largest coefficient of variation) at the border of the coexistence region where it either appears or disappears (Fig. 2c). As a species reaches the point where it is no longer able to persist in the community, both its mean biomass and its variance decrease linearly to zero, so that its invariability (the ratio of squared mean to variance) also decreases linearly to zero. Thus, on its path to disappearance, the rarer the species, the more variable it is. Eventually, at the point of disappearance, the invariability of this species drops to zero, just like asymptotic resilience.

As a consequence, asymptotic resilience *ℜ* and minimum invariability ℐ_min_ display the same stability patterns (compare Figs. 2e and 2f). Recall that minimum invariability is defined as the invariability of the most variable population, that is, the minimum of the curves in Fig. 2c. A species that is either close to being excluded from the community or close to being able to invade the community governs the dominant eigenvalue and is also the most variable species. Both the dominant eigenvalue and that species’ invariability become zero at its extinction/invasion threshold, so that asymptotic resilience and minimum invariability have the same behavior close to this threshold. Moreover, because these thresholds occupy the entire gradient of resource supply, the two stability measures show similar patterns along the gradient. This similarity suggests that asymptotic resilience is effectively an extreme version of population-level stability, determined by the most unstable species, which is almost always rare.

Population invariability ℐ_pop_ combines the invariabilities of individual populations in a more balanced way (Fig. 2g). In the regions where a single species dominates, it is equal to the invariability of the dominant species. In the coexistence regions, it switches smoothly from the invariability of the species that dominates at one border to the invariability of the species that dominates at the other border. As a result, the large variability (invariability dropping to zero) of the species that becomes rare in the coexistence region is not visible in the pattern of population invariability ℐ_pop_. This is in sharp contrast to the patterns of asymptotic resilience *ℜ* and minimum invariability ℐ_min_, which drop to zero each time the invariability of an individual population drops to zero.

Ecosystem invariability ℐ_eco_ quantifies ecosystem-level stability. While mean total biomass is approximately constant along the gradient, the variance of total biomass decreases in the coexistence regions (Fig. 2d), so that ecosystem invariability ℐ_eco_ increases (Fig. 2h). Hence, the effect of coexistence on ℐ_eco_ is opposite to its effect on ℐ_pop_. Population invariability ℐ_pop_ decreases in the coexistence regions, because the coexisting species have smaller mean biomass and larger variance compared to the regions where they dominate. In contrast, ecosystem invariability ℐ_eco_ increases, because species fluctuations partially compensate each other. Note that both measures ℐ_pop_ and ℐ_eco_ depend on common species and change smoothly along the gradient of resource supply.

**Figure 2:**
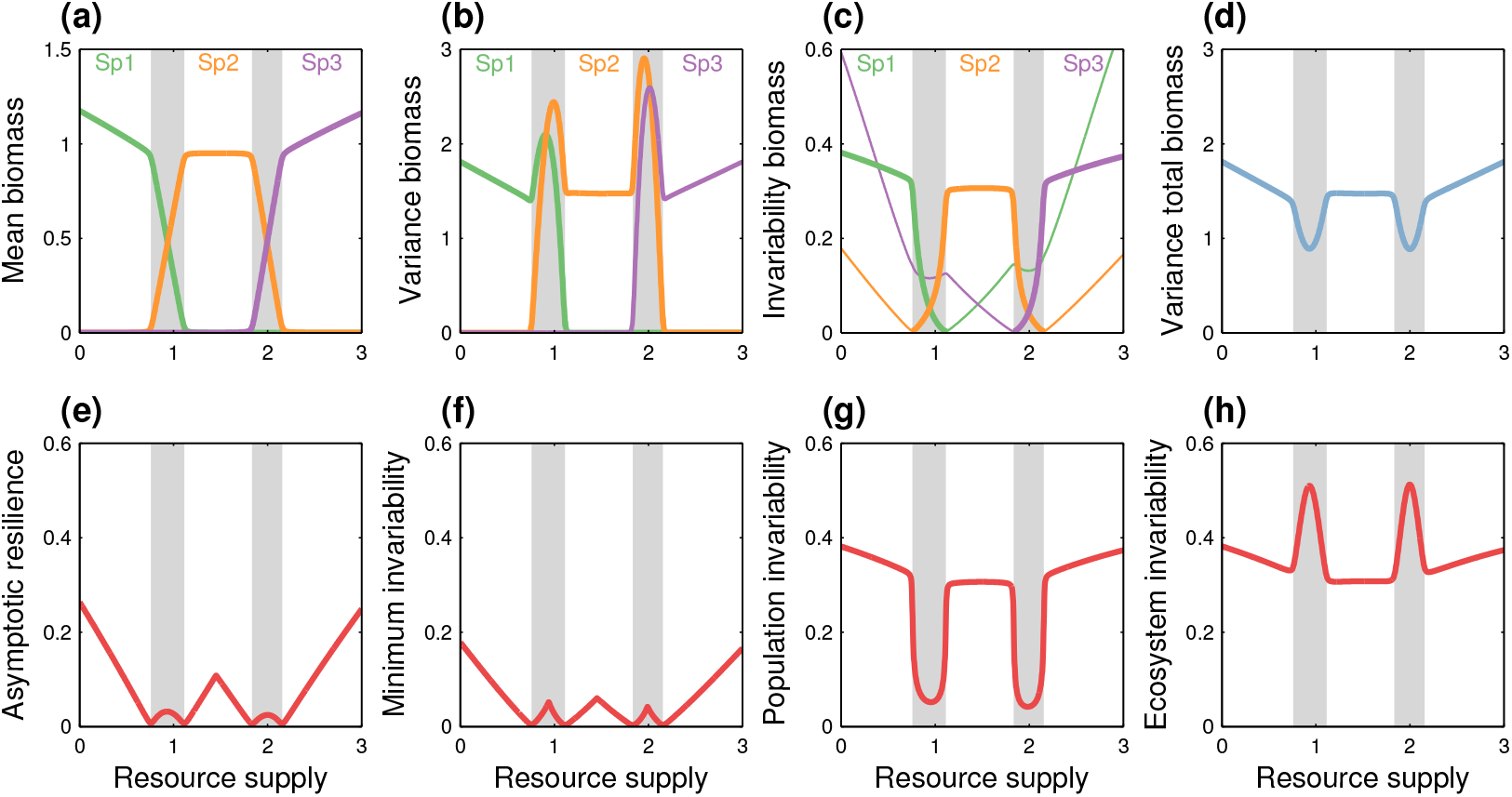
Stability in a resource competition model. (a) Community composition changes along a gradient of resource supply. Coexistence regions are indicated in grey. Note that mean total biomass is roughly constant. (b) When applying a white-noise perturbation, the biomass of individual populations has larger variance in the coexistence regions than in the regions where a single species dominates. (c) The invariability of a population is smallest at the border of the coexistence regions where it is outcompeted by another species. Thin lines represent rare populations that are maintained due to immigration (sink populations; see also Fig. S3). (d) The variance of total biomass is smallest in the coexistence regions. (e) Asymptotic resilience *ℜ* _drops_ to zero at the borders of the coexistence regions. (f) Minimum variability ℐ_min_ has a behavior similar to asymptotic resilience. (g) Population invariability ℐ_pop_ is smallest in the coexistence regions. (h) Ecosystem invariability ℐ_eco_ is largest in the coexistence regions.

## Consistency across levels of organization

The previous section shows that, for a model formulated at the population level, asymptotic resilience is an extreme version of population-level stability. This suggests that asymptotic resilience might be used as a measure of stability at the ecosystem level by constructing a model with aggregated ecosystem-level variables. But it is unclear whether asymptotic resilience in the aggregated ecosystem-level model will bear any relationship to asymptotic resilience in the population-level model. Here we examine the consistency of asymptotic resilience and invariability as measures of stability as the system is scaled up from the population to the ecosystem level.

To construct an upscaled model of the consumer-resource system, we lump the three consumer biomasses into a single dynamical variable, which can be interpreted as the biomass of an aggregated consumer that combines the features of the three consumer species (see Appendix S2 for details). The aggregated consumer uses the resources in the same way as the species that is most efficient under the prevailing environmental conditions, and thus it switches between the consumption characteristics of the various species along the gradient.

Asymptotic resilience yields results that differ both qualitatively and quantitatively between the population- and ecosystem-level models (Fig. 3a). In particular, the stability loss associated with each extinction or invasion event in the population-level model is absent from the ecosystem-level model. Moreover, the pattern in the ecosystem-level model is difficult to interpret. For example, asymptotic resilience predicts a sharp stability increase close to the borders of the environmental gradient, but it is unclear what mechanism causes these two peaks. In contrast, ecosystem invariability ℐ_eco_ has a consistent behavior between the two models (Fig. 3b). The stability pattern in the ecosystem-level model is similar, both qualitatively and quantitavely, to that of the population-level model, except that the humps in the coexistence regions have disappeared. Clearly, this is due to the fact that the ecosystem-level model does not contain the details of individual consumer species and their coexistence.

Thus, ecosystem invariability has a consistent and predictable behavior in the model upscaling, while the pattern predicted by asymptotic resilience undergoes drastic changes with upscaling.

## Diversity-stability relationships

We build now upon the results of the previous sections to connect different predictions about the diversity-stability relationship. By taking into account the level of organization addressed by the various stability measures, we obtain a framework that coherently integrates the different theoretical predictions and that is consistent with empirical data.

Random interaction models have been widely used in theoretical studies of ecological stability since May’s (1973) seminal work. Most of these studies generate random community matrices and study their stability properties, e.g., the proportion of stable matrices, or the asymptotic resilience *ℜ* of those matrices that are stable. However, the invariability measures we have introduced also depend on the equilibrium abundance vector *N^∗^*. Therefore, we take a step back from the standard approach, and start by generating random interaction matrices. We then look for the equilibrium to which the community dynamics converge. This ensures that the equilibrium, in which some species may be absent, is stable. We then quantify the stability of this equilibrium, that is, we determine the equilibrium vector *N^∗^* and the community matrix *A*, and compute the corresponding stability measures *ℜ*, ℐ_min_, ℐ_pop_ and ℐ_eco_. We consider a competitive Lotka-Volterra model in which the interspecific competition coefficients are randomly drawn and mutually independent (see Appendix S4 for details). Fig. 4 shows the dependence of the four stability measures on the number of species present at equilibrium (the results are qualitatively the same for the number of species in the initial species pool, see Fig. S4).

**Figure 3:**
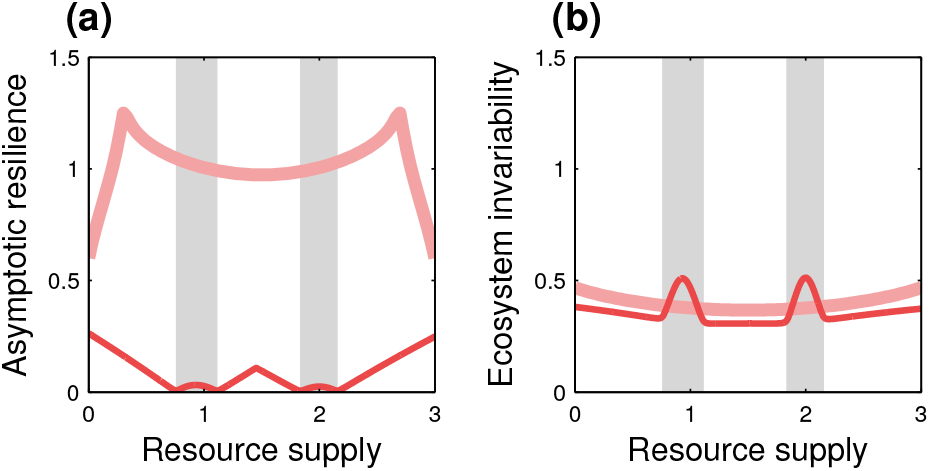
Are stability measures consistent across levels of organization? Consumer species in the resource competition model (Fig. 2) are aggregated into a single consumer variable. (a) Asymptotic resilience *ℜ* changes dramatically between the population-level (dark thin line) and ecosystem-level (light thick line) models. (b) Ecosystem invariability ℐ_eco_ has a similar behavior in the population-level and ecosystem-level models, although some details related to species coexistence are lost during model reduction.

The three stability measures that address population-level stability (i.e., *ℜ*, ℐ_min_ and ℐ_pop_) predict negative relationships between diversity and stability, while ecosystem invariability predicts a positive relationship. This shows that the diversity-stability relationship is strongly dependent on the level of organization addressed by the measure used to quantify stability. These contrasting patterns have been observed in other competition models (May, 1974; Lehman & Tilman, 2000; Loreau & de Mazancourt, 2008) and are consistent with empirical data (Tilman *et al.*, 2006; Jiang & Pu, 2009; Campbell *et al.*, 2011; Gross *et al.*, 2014).

Furthermore, asymptotic resilience *ℜ* and minimum invariability ℐ_min_ display very similar relationships, in agreement with the results of our resource competition model. Compared with population invariability ℐ_pop_, the dependence of *ℜ* and ℐ_min_ on species richness is stronger and has a larger dispersion. This is consistent with the fact that *ℜ* and ℐ_min_ are extreme measures of population-level stability that drop to zero at the borders of the stability regions (see Fig. 2).

## Discussion

Resilience and variability are the most widely used notions of ecological stability in theoretical and empirical studies, respectively. We have shown that they have fundamentally different properties. The stability patterns across ecosystems predicted by asymptotic resilience *ℜ* are largely determined by changes in the equilibrium community composition (i.e., the set of persistent species). Because asymptotic resilience drops to zero at each compositional change, its patterns consist of a sequence of humps in between the zeros, each hump zooming in on a specific community. In contrast, population invariability ℐ_pop_ and ecosystem invariability ℐ_eco_ are only weakly sensitive to compositional changes, so that their patterns connect smoothly communities with different composition.

**Figure 4:**
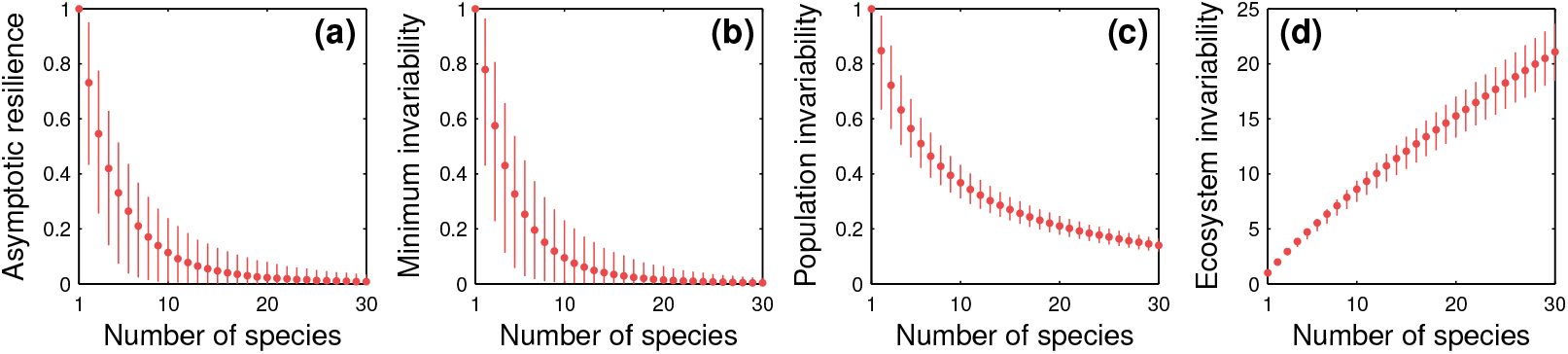
Stability in a competitive Lotka-Volterra model with random interactions. Dots and vertical lines show the median and the range from the 5th to the 95th percentile, respectively. With increasing realized species richness, *ℜ*, ℐ_min_ and ℐ_pop_ decrease, while ℐ_eco_ increases. Note the relatively large dispersion of *ℜ* and ℐ_min_, and the smaller dispersion of ℐ_pop_ and ℐ_eco_.

To better understand the differences between resilience and invariability, we have introduced a very particular invariability measure, minimum invariability ℐ_min_, or the invariability of the most variable species, which displays patterns similar to asymptotic resilience. Both measures drop to zero at each species gain or loss, so that the similarity is particularly strong when compositional changes occur repeatedly, e.g., along an environmental gradient. Although this similarity is not a mathematical equivalence, it strongly suggests that asymptotic resilience is a stability measure that generally focuses on the most variable species.

We have also shown that rare species strongly affect these two measures. Although not every rare species is highly variable, the most variable species in a community is nearly always rare. For example, a species that is present in a community due to immigration without being able to maintain itself (a sink population) is typically rare but not necessarily variable (i.e., it can have a small coefficient of variation; see thin lines in Fig. 2c). Its variability is high only when it gets close to its invasion threshold. Note that in real-world communities many rare species are probably occasional immigrants that do not take part in community dynamics (even if they are close to invasion). In this case, the stability information provided by asymptotic resilience is of little relevance from an ecosystem perspective.

The family of invariability measures is more flexible than asymptotic resilience. Measures ℐ_pop_ and ℐ_eco_ can be used to consistently quantify stability not only at different levels of organization within the same model, but also across models formulated at different levels of organization. Population and ecosystem invariabilities, unlike asymptotic resilience, do not drop to zero when species composition changes. This might seem surprising, as zero asymptotic resilience indicates an instability in the system. To understand this, it is important to recall that the stochastic perturbation inducing system variability acts on a per capita basis in our model. When a species approaches its extinction or invasion threshold, the perturbation applied to this species tends to zero, so that the corresponding instability is less and less excited. As a result, this instability does not show up in the patterns of population and ecosystem invariabilities (but it does in the pattern of minimum invariability). In contrast, due to its rigidity, asymptotic resilience is insensitive to the magnitude and direction of the perturbation.

Less extreme versions of asymptotic resilience exist, such as non-asymptotic return rates (the rates at which a system returns to equilibrium, measured at finite times after a pulse perturbation) and return times (the times it takes after a pulse perturbation for the system to return to, and remain within, certain distances from the equilibrium). These alternative measures, which have been used in empirical studies (e.g., Steiner *et al.*, 2006; Sibly *et al.*, 2007), are promising from a theoretical viewpoint. Because variability can be seen as a superposition of short- and long-term responses to pulse perturbations (see Introduction), one can expect that short-term return rates should have similar properties as the invariability measures we have studied. In particular, short-term return rates do not have the rigidity of asymptotic resilience. Further work in this direction could enable a closer interaction between theoretical and empirical studies on the response of ecosystems to pulse perturbations.

The various stability measures we have discussed rest on the equilibrium assumption. Obviously, this assumption is ingrained in resilience measures based on the return to equilibrium. In theory, one could extend our analysis to non-equilibrium unperturbed states (e.g., limit cycles). For variability-based measures, the equilibrium assumption excludes the possiblity that the dynamics generate variability autonomously (e.g., limit cycles or more complex attractors). Several theoretical stability studies have quantified stability based on the intensity of internally generated variability (e.g., McCann *et al.*, 1998; Brose *et al.*, 2006). It is unclear whether the latter variability is related to the externally generated variability we have studied. While there is some empirical support for internally generated variability, we believe that externally generated variability continues to deserve research attention, especially because it is likely more common and it allows the elaboration of a more systematic theory.

Our findings have some bearing on the theory of tipping points (Scheffer *et al.*, 2009). First, it is important to note that the extinction/invasion bifurcations we have analyzed are of a different type than the bifurcations studied in the theory of tipping points. While in the former bifurcations the equilibrium changes smoothly (the biomass of a species decreases to zero or increases from zero), the latter bifurcations are characterized by sudden shifts from one equilibrium to another dynamical state. The theory of tipping points is mainly concerned with the properties of the equilibrium when approaching the bifurcation, and in particular, whether the upcoming transition can be detected beforehand (early warning signals). Usually this detection is based on increased variability: when approaching the bifurcation, stability is gradually lost (asymptotic resilience decreases to zero) and variability diverges. Our work implies that this detection mechanism might fail for extinction/invasion bifurcations. When approaching such a bifurcation, stability is gradually lost, but the variability of most system variables does not diverge. This illustrates that bifurcations are not necessarily accompanied by early warning signals.

Lastly, we have applied these insights to the interpretation of diversity-stability relationships. The random interaction model we have used for this purpose differs from previous such models, which quantified the probability to encounter stable community matrices (May, 1973; Allesina & Tang, 2013; Coyte *et al.*, 2015). Instead, we studied the stability properties of communities that are realized (and therefore stable) starting from a larger species pool. We found consistent relationships depending on the level of organization at which stability is quantified. Ecosystem-level stability tends to increase with diversity, while population-level stability tends to decrease with diversity. These predictions agree well with observations from grasslands (Gross *et al.*, 2014) and other competitive systems (Jiang & Pu, 2009; Campbell *et al.*, 2011). It would be interesting to extend this comparison to multi-trophic systems, for which the empirical patterns are less clear-cut (Jiang & Pu, 2009; Campbell *et al.*, 2011). The theoretical study of diversity-stability relationships in food webs presents new challenges, as the predictions of random interaction models are known to be sensitive to specific structural constraints (Neutel *et al.*, 2002; Rooney *et al.*, 2006; Brose *et al.*, 2006; Allesina *et al.*, 2015).

By showing that asymptotic resilience behaves like the invariability of the most variable species, our work translates results obtained in the theoretical literature to the empirical world. By developing theoretical measures of invariability, our work paves the way to further develop theory in touch with the empirical world. Further studies might be able to extend our work to non-equilibrium conditions, or integrate other important stability measures, such as resistance to perturbations or return times to equilibrium. Meanwhile, the general relationships between stability measures we have uncovered should be instrumental in integrating empirical and theoretical approaches to ecological stability.

## Acknowledgements

This work was supported by the TULIP Laboratory of Excellence (ANR-10-LABX-41 and ANR-11-IDEX-002-02), by a Region Midi-Pyrénées project (CNRS 121090), and by the BIOSTASES Advanced Grant, funded by the European Research Council under the European Union’s Horizon 2020 research and innovation programme (grant agreement number 666971).

